# Explicit Scale Simulation for analysis of RNA-sequencing with ALDEx2

**DOI:** 10.1101/2023.10.21.563431

**Authors:** Gregory B. Gloor, Michelle Pistner Nixon, Justin D. Silverman

## Abstract

In high-throughput sequencing (HTS) studies, sample-to-sample variation in sequencing depth is driven by technical factors, and not by variation in the scale (e.g., total size, microbial load, or total mRNA expression) of the underlying biological systems. Typically a statistical normalization is used to remove unwanted technical variation in the data or the parameters of the model to enable analyses that are reliant on scale; e.g., differential abundance and differential expression analyses. We recently showed that all normalizations make implicit assumptions about the unmeasured system scale and that errors in these assumptions can dramatically increase false positive and false negative rates. We demonstrated that these errors can be mitigated by accounting for uncertainty about scale using a *scale model*, which we integrated into the ALDEx2 R package. This article provides new insights into those methods, focusing on the application to transcriptomic analysis. Here we provide transcriptomic case studies demonstrating how scale models, rather than traditional normalizations, can reduce false positive and false negative rates in practice while enhancing the transparency and reproducibility of analyses. We show that these scale models replace the need for dual cutoff approaches often used to address the disconnect between practical and statistical significance. We demonstrate the utility of that scale models built based on known housekeeping genes in complex metatranscriptomic datasets. Thus this work provides example and practical guidance on how to incorporate scale into transcriptomic analysis.

## Introduction

High-throughput sequencing (HTS) is a ubiquitous tool used to explore many biological phenomenon such as gene expression (single-cell sequencing, RNA-sequencing, meta-transcriptomics), microbial community composition (16S rRNA gene sequencing, shotgun metagenomics) and differential enzyme activity (selex, CRISPR killing). HTS proceeds by taking a sample from the environment, making a library, multiplexing (merging) multiple libraries together, and then applying a sample of the multiplexed library to a flow cell. Each of these steps is a compositional sampling step as only a fixed-size subsample of nucleic acid is carried over to subsequent steps. Thus, with each sampling step the connection between the actual size of the sampled DNA pool and the scale (e.g., size, microbial load, or total gene expression) of the measured biological system is degraded or lost. In the end, the information contained in the data relates only to relative abundances and has an arbitrary scale imposed by the sequencing process (1–3). Researchers can use modified experimental protocols (4–7) or machine learning methods (8) to uncover the biological variation in scale. However, wet-lab protocols only provide information on the size of the data downstream of the step in the sample preparation protocol where the intervention was made and introduce an additional source of variation that must be accounted for (5). The computational methods developed for microbiome analysis have low correlation with the actual scale but are useful (8). In short, the disconnect between sample-to-sample variation in sequencing depth and biological variation in scale remains an open challenge.

The analysis of HTS data suffers from several known problems that can be traced, in whole or in part, to misspecification of scale. The first issue is poor control of the false discovery rate (FDR) (9–13), exhibited as dataset-dependent FDR control and by the disconnect between statistical and biological significance (14). In current practice, these issues are addressed by a dual-filtering method, whereby both a low p-value (or equivalently a low q-value following FDR correction (15)) and a large difference between groups is used to identify interesting transcripts or genes for follow-up analysis (14, 16). This double-filtering approach is graphically exemplified by the volcano plot (16), but is known to not appropriately control the FDR (17, 18). The second issue is poor performance when analyzing data where the mean change between groups is non-zero (3). Such asymmetric data can arise when a gene set is expressed in one group but not the other, or when one group contains different gene content from the other. This type of data frequently arises in in-vitro selection experiments (SELEX), transcriptome analysis, and microbiome analysis (19). The third issue is that the actual scale of the environment is often a major confounding variable during analysis (3, 8). The final issue is that these problems become more pronounced as more samples are collected; that is, more information results in a worsening of the accuracy of the analysis (3, 13, 20).

The four problems were recently shown by Nixon et al. (3) to be a result of a mismatch between the underlying size or scale of the system and the assumptions of the normalizations used for the analysis of HTS. Biological variation in scale often represents an important unmeasured confounder in HTS analyses (21). For example, cells transformed by the cMyc oncogene have about 3 times the amount of mRNA and about twice the rRNA content than non-transformed cells (22), and this dramatically skews transcriptome analysis unless spike-in approaches are used (5). In addition, wild-type and mutant strains of cell lines, yeast or bacteria have different growth rates and RNA contents under different conditions, which affect our ability to identify truly differentially abundant genes (23–25). As another example, the total bacterial load of the vaginal microbiome differs by 1-2 orders of magnitude in absolute abundance between the healthy and bacterial vaginosis states (26), and the composition between these states is dramatically different (27, 28). Thus, a full description of any of these systems includes both relative change (composition) and absolute abundance (scale). Current methods access only the compositional information yet make implicit assumptions about the scale (20).

Recently, Nixon et al. (3) showed that the challenge of non-biological variation in sequencing depth be viewed as a problem of partially-identified models. They showed that *all* normalizations make some assumption about scale but these implicit assumptions are often inappropriate and difficult to interpret. This causes different normalizations to provide different outputs when applied to the same dataset (9, 11, 29–31). Intuitively, normalizations in widespread use assume that either all samples have the same scale, e.g. proportions, rarefaction (32), RPKM (33, 34), etc; or that a subset of features in one sample can be chosen as a reference to which the others are scaled e.g. the TMM (35), or LVHA (19) or the additive log-ratio (36); or that different sub-parts of each sample maintain a constant scale across samples e.g. the RLE (37); or that the geometric mean of the parts is appropriate e.g the CLR (38) and its derivatives.

The original naive ALDEx2 (39) model unwittingly made a strict assumption about scale through the CLR normalization (3). This assumption was often close enough to the true value to be useful, but was not always the a good estimate and could be outperformed by other normalizations (40). Nixon et al. (3) showed that better scale assumptions resulted in more reproducible data analysis including better control of both false positive and false negative results. We recently modified ALDEx2 to explicitly model the scale over a range of reasonable normalization parameters, and showed significant improvements in performance in microbiome and in-vitro selection experiments (20). Here, we briefly review these modifications and show how scale uncertainty can greatly improve modeling in transcriptome and meta-transcriptome datasets to provide more robust and reproducible results.

### Implementation

Formal and expanded descriptions of the concepts that follow are given in (3, 20). To be concrete, we let **Y** denote the *measured D × N* matrix of sequence counts with elements **Y**_*dn*_ indicating the number of measured DNA molecules mapping to feature *d* (e.g., a taxon, transcript or gene) in sample *n*. Likewise, we can denote **W** as the *true* amount of class *d* in the biological system from which sample *n* was obtained. We can think of **W** as consisting of two parts, the scale **W**^⊥^ (e.g., totals) and the composition **W**^∥^ (i.e., proportions).

That is, **W**^⊥^ is a *N* -vector with elements 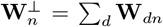 while **W**^∥^ is a *D × N* matrix with elements 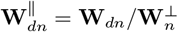. Note that with these definitions **W** can be written as the element-wise combination of scale and composition: 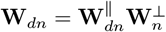, or as the logarithm log 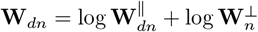.

Many of the normalizations in widespread use in tools such as DESeq2 (41), edgeR (35), metagenomeSeq (42) ALDEx2 (43) can be stated as ratios of the form **ŵ** _*dn*_ ≈ **Y**_*dn*_*/f* (**Y**), where the denominator is determined by some function of the observation. We use the **ŵ** (^) notation to indicate that the output is an estimate of the true value. The technical variation in sequencing depth 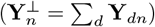 implies that observed data **Y** provides us with information about the system composition **W**^∥^ but little to no information in the system scale **W**^⊥^ (Lovell et al. 2011).

### Adding Scale Uncertainty in ALDEx2

The ALDEx2 R package (39, 43) is a general purpose toolbox to model the uncertainty of HTS data and to use that model to estimate the underlying LFC (log-fold change) significance. At a high-level, ALDEx2 has three connected components to estimate the uncertainty inherent in HTS datasets. First, the tool accounts for the uncertainty of the sequencing counts using Dirichlet multinomial sampling to build a probabilistic model of the data; i.e., **ŵ** ^∥^ ≈ Dir(**Y**). Secondly, ALDEx2 uses the centred log-ratio transformation to scale the data (39). However, this step was modified recently to account for scale uncertainty and misspecification

(20) via a scale model, explained with more details in (3, 20) and summarized in the next paragraph. Finally, a standard null-hypothesis test and a non-parametric estimate of mean standardized difference are used to report on the finite sample variation. These sources of uncertainty and variation are combined via reporting the expected values from a Monte-Carlo simulation framework. For simplicity, we use the term ‘difference’ to refer to the absolute difference between groups, and ‘dispersion’ to refer to the within-condition difference or pooled variance as defined in (39). These are calculated on a log_2_ scale. For more details on ALDEx2 see (3, 20, 39, 43).

Scale models can be incorporated into ALDEx2, turning the ALDEx2 model into a specialized type of statistical model which Nixon et al. (3) term a *Scale Simulation Random Variable* (SSRV). To do this, Nixon et al. (3) generalized the concept of normalizations by introducing the concept of a *scale model* to account for potential error in the centred log-ratio normalization step. They did this by including a model for 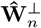. The CLR normalization used by ALDEx2 makes the assumption 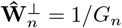, where *G*_*n*_ is the geometric mean of sample n, which while being a random variable, is essentially constant across each Monte-Carlo replicate, but that differs between samples. With this modification, ALDEx2 can be generalized by considering probability models for the scale 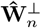 that have mean 1*/G*_*n*_. For example, the following scale model generalizes the CLR:

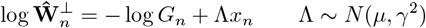

This formulation is quite flexible (3, 20). In the simple or ‘default’ configuration, *µ* = 0 and *γ* is a tunable parameter drawn from a log-Normal distribution(3). Adding scale uncertainty with the *γ* paramenter controls only the degree of uncertainty of the CLR assumption for the *x*_*n*_ binary condition indicator (e.g., *x*_*n*_ = 1 denotes case and *x*_*n*_ = 0 denotes control). In the advanced or ‘informed’ configuration, *µ* takes different values for each group and controls the location of the LFC assumption; combining *µ* with a *γ* estimate allows for uncertainty in both the location and the scale. An example of both the default and informed approaches is given for a microbiome dataset in (20) showing increased sensitivity and specificity. Here we show that these approaches also work well in transcriptome and metatranscriptome datasets. These modifications are instantiated in ALDEx2 which is the first software package designed for SSRV-based inference.

## Results

### Adding scale uncertainty replaces the need for dual significance cutoffs

Gierliński et al. (44) generated a highly replicated yeast transcriptome dataset to compare gene expression between a wild-type strain and a snf2 gene knockout, Δsnf2. This dataset was used to test several RNA-seq tools for their power to detect the set of differentially abundant transcripts identified in the full dataset when the data was subset (14). In this study each tool had its own ‘gold standard’ set of transcripts with different tools identifying between between 65% to >80% of all transcripts as being significantly different. Since the majority of transcripts were significantly different, the authors suggested that it was more appropriate to apply a dual cutoff composed of both a Benjamini-Hochberg (45) corrected p-value (q-value) plus a difference cutoff to limit the number of identified transcripts to a much smaller fraction of the total. Nixon et al, (20) showed that adding even a small amount of scale uncertainty with ALDEx2 dramatically reduced the number of significant transcripts identified, removing the need for the dual cutoff approach in this dataset and others. Below we include an intuitive explanation of why and how incorporating scale uncertainty achieved this outcome using a setting of *γ* = 0.5. The approach is in line with the recommendations of (20) and gives results comparable to those proposed in (14).

We start with the assumption that not all statistically significant differences are biologically relevant (46), and that such a large number of significantly different transcripts breaks the necessary assumption for DA/DE expression that most parts be invariant (30). As noted, transcriptomics commonly uses a dual cutoff approach that is graphically exemplified by volcano plots (14, 16). Using either DESeq2 or ALDEx2, a majority of transcripts are statistically significantly different between groups with a q-value cutoff of ≤ 0.05; i.e. 4636 (79%, DESeq2) or 4172 (71%, ALDEx2) of the 5891 transcripts. These values are in line with those observed by (14). Such large numbers of statistically significant transcripts seems biologically unrealistic. That 118 transcripts are identified by ALDEx2 and not DESeq2, while DESeq2 identifies 582 transcripts that ALDEx2 does not, suggests that the choice of normalization plays a role in which results are returned as significant and that some, if not the majority, are driven by technical differences in the analysis (13, 30).

The Volcano plots in Figure 1 A and B show that adding scale uncertainty increases the minimum q-value and increases the concordance between the q-value and the difference between groups (compare panels A and B). The effect plots (47) in Figure 1C shows that the majority of significant transcripts (red, orange) have negligible differences between groups and very low dispersion. We suggest that this low dispersion is driven by the experimental design which is actually a technical wet lab replication rather than a true biological replication design (44). Scale uncertainty can be incorporated using the `gamma` parameter that controls the amount of noise added to the CLR mean assumption when we call either `aldex()`, or `aldex.clr()`. Figure 1 B,D shows that setting *γ* = 0.5 results in 205 which is far fewer significant transcripts than in the naive analysis and we observe that the minimum dispersion increases from 0.12 (*γ* = 0) to 0.67 (*γ* = 0.5).

**Figure 1:**
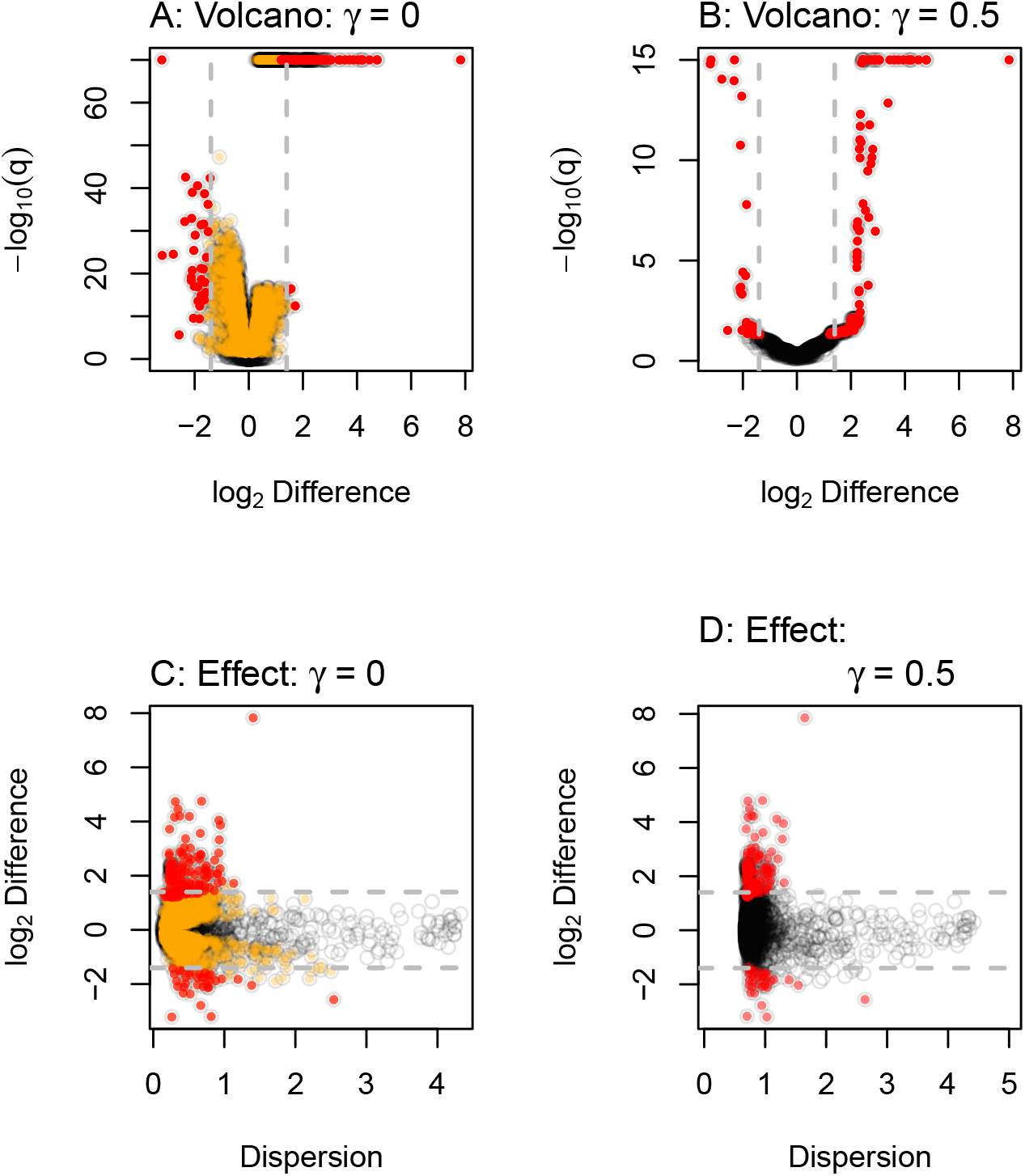
Volcano and effect plots for unscaled and scaled transcriptome analysis. ALDEx2 was used to conduct a differential expression (DE) analysis on the yeast transcriptome dataset. The results were plotted to show the relationship between difference and dispersion using effect plots or difference and the q-values using volcano plots. Panels A,C are for the naive analyses, and Panels B,D are for the default analyses that include scale uncertainty. Each point represents the values for one transcript, with the color indicating if that transcript was significant in the both analyses (red) or in the naive analysis only (orange). Points in grey are not statistically signficantly different under any condition. The horizontal dashed lines represent a log_2_difference of ±1.4.

It is common practice to use a dual-cutoff by choosing transcripts based on a thresholds for both q-values and fold-changes (14, 16). Note that there is considerable variation in recommended cutoff values(14). Here, applying a dual-cutoff using a heuristic of at least a 2^1.4^ fold change reduces the number of significant outputs to 193 for DESeq2 and to 186 for ALDEx2. This cutoff was chosen for convenience and is in-line with the recommendations of (14) with the fold change limits shown by the dashed grey lines in Figure 1. The 2^1.4^ fold change cutoff identifies a similar number of transcripts as does ALDEx2 using *γ* = 0.5 which identifies 205. Supplementary Figures 1 and 2 shows how to use the `aldex.senAnalysis()` function to identify those transcripts that are very sensitive to scale uncertainty. In this supplementary figure we see that even adding a very small amount of scale *γ* = 0.1 reduces the number of significant transcripts by more than half. This allows us to ignore those low-dispersion transcripts that were significant only because of an absence of scale uncertainty. In practice, we suggest that a `gamma` parameter between 0.5 and 1 is realistic for most experimental designs (20).

**Figure 2:**
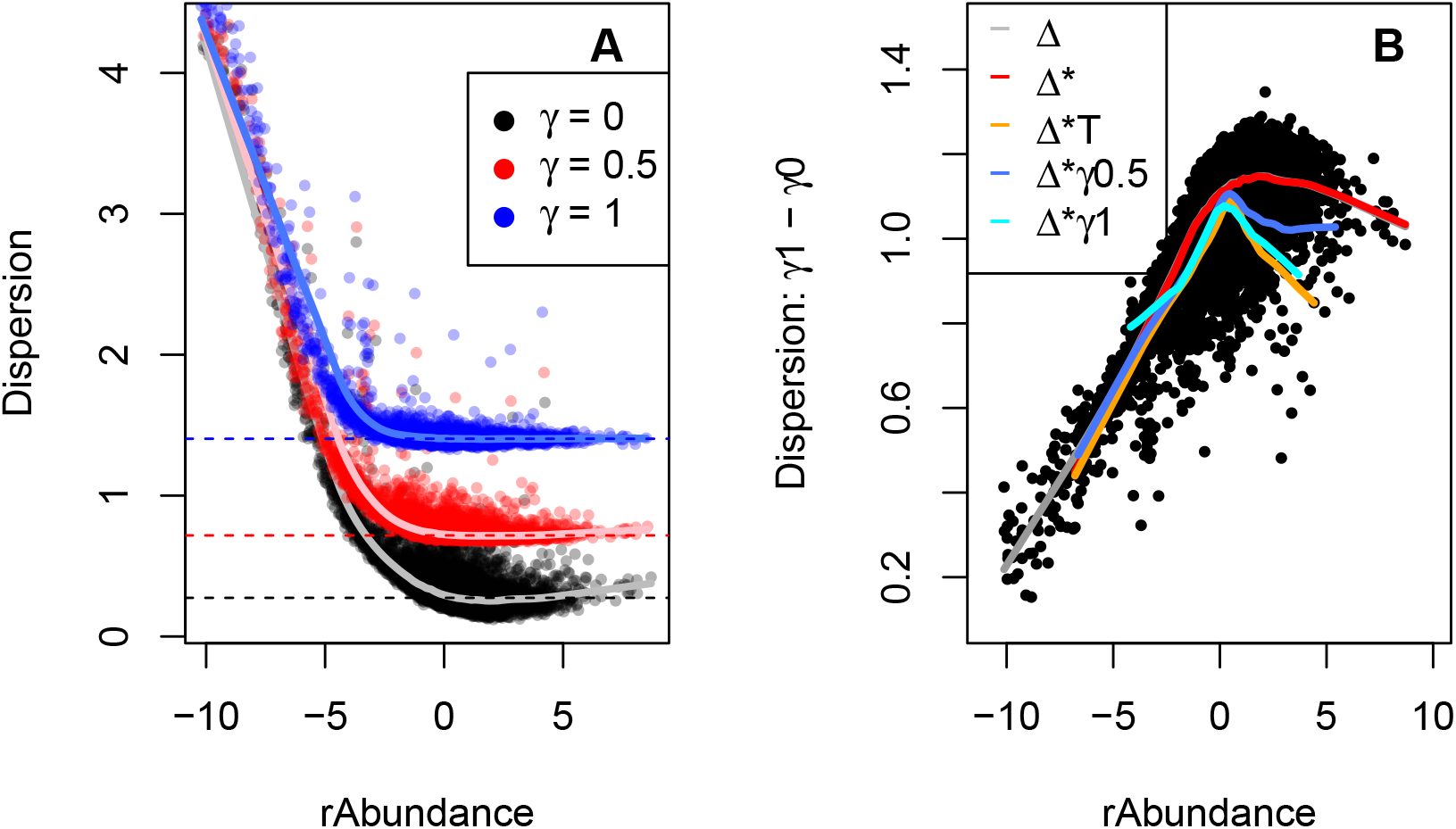
Adding scale uncertainty changes the dispersion distribution. Panel A shows a plot of the expected value for relative abundance vs the expected value for the pooled dispersion as output by aldex.effect. The dashed horizontal lines show the median value for the features with a rAbundance between -0.5 and 0.5, and the light colored lines are lowess lines of fit through the center of mass of the data. Panel B plots the dispersion difference between *γ* = 1 and *γ* = 0; note the non-linear relationship that highlights the rotation that is evident in Panel A. The colored lines indicate the lowess line of fit through the centre of mass of the plot for the various populations of points. The grey line is the total population and shows the difference Δ, the red line is the population of significant transcripts (*) with *γ* = 0, the orange line is the population of significant transcripts with a difference threshold (T) of about ±2^1.4^-fold change, the blue line is the population of significant transcripts with *γ* = 0.5, and the cyan line is the significant population with *γ* = 1. Δ: Difference, *: significant, T: thresholded.

The effect on dispersion with increasing amounts of scale uncertainty are shown in Figure 2A, where we can see that the dispersion increases as uncertainty is added. Note that the dispersion in the unscaled analysis in Figure 2A reaches a minimum near the mid-point of the distribution, and also does so when the analysis is conducted with DESeq2 (Supplementary Figure 3). This shows more clearly that dispersion of many transcripts is almost negligible in the absence of scale uncertainty. This plot makes the counter-intuitive suggestion that the variance in expression of the majority of genes with moderate expression is more predictable than highly-expressed genes or of housekeeping genes (48). This is at odds with the known biology of cells where single cell counting of highly-expressed transcripts shows that they have little intrinsic variation (23, 49).

Adding scale uncertainty by setting *γ* = 0.5, or *γ* = 1.0, increases the minimum dispersion as shown in Figure 2A by the red and blue data points, and by the colored lines of fit through the centre of mass of the data. Less obvious is that the additional dispersion is not applied equally to all points. Figure 2B shows a plot of the difference between the *γ* = 0 and *γ* = 1 data and here we can see that scale uncertainty is preferentially increasing the dispersion of the mid-expressed transcripts that formerly had negligible dispersion; examine the grey line of best fit (overlaid by the red line) for the trend. Panel B also shows the trend of the expression-dispersion relationship for transcripts that are classed as statistically significant. The red line shows the trendline with no added scale uncertainty, and this trendline exactly overlays with the grey trendline for the bulk of transcripts. The orange trendline indicates those transcripts that are both statistically significant and that have a thresholded expression level of ±1.4, and the dark blue and cyan lines show the statistically significant trendline for *γ* = 0.5, or 1.0.

Thus, taking Figures 1 and 2 together adding scale uncertainty has the desirable effect of changing the distribution of transcripts identified as significantly different between groups. Those parts that were statistically significantly different *only because of low dispersion* are preferentially excluded from statistical significance while those parts that were significantly different because of a high difference between groups remain.

### Housekeeping genes and functions to guide scale model choices

Dos Santos et al. (50) used a vaginal metatranscriptome dataset to compare the gene expression in bacteria collected from healthy (H) and bacterial vaginosis (BV) affected women. In this environment, both the relative abundance of species between groups and the gene expression level within a species is different (51). Additionally, prior research suggests that the total number of bacteria is about 10 times more in the BV than in the H condition (26). Thus, these are extremely challenging datasets in which to determine differential abundance as there are both compositional and scale changes between conditions. The usual method to analyze vaginal metratranscriptome data is to do so on an organism-by-organism basis (51–53) because the scale confounding of the environment is less pronounced. One attempt at system-wide analysis returned several housekeeping functions as differentially expressed between groups (52); a result likely due to a disconnect between the assumptions of the normalization used and the actual scale of the environment (19).

In this example, we show how to specify and interpret an informed scale model that can explicitly account for some of these modeling difficulties (20) even in a difficult dataset. An informed scale model can control for both the mean difference of scale between groups (e.g., directly incorporate information on the differences in total number of bacteria between the BV and H conditions) as well as the uncertainty of that difference. To specify a user-defined scale model, we can pass a matrix of scale values instead of an estimate of just the scale uncertainty to `aldex.clr()`. This matrix should have the same number of rows as the of Monte-Carlo Dirichlet samples, and the same number of columns as the number of samples. While this matrix can be computed from scratch by the analyst, there is an `aldex.makeScaleModel()` function that can be used to simplify this step in most cases. This encodes the scale model as Λ ∼ *N* (*log*_2_*µ*_*n*_, *γ*^2^), where *µ*_*n*_ represents the scale value for each sample or group and gamma is the uncertainty as before. The scale estimate can be a measured value (cell count, nucleic acid input, etc) or an estimate. Nixon et al. (3, 20) showed that only the ratio of the means are important when providing values for *µ*_*n*_; i.e., the ratio between the log_2_ *µ*_*i*_ and log_2_ *µ*_*j*_ values. See the supplement to Nixon et al. (20) for more information.

Figure 3A shows an effect plot of the data where reads are grouped by homologous function regardless of the organism of origin. Each point represents one of 3728 KEGG functions (54). There are many more functions represented in the BV group (bottom) than in the healthy group (top). This is because the *Lactobacilli* that dominate a healthy vaginal microbiome have reduced genome content relative to the anaerobic organisms that dominate in BV, because there is a greater diversity of organisms in BV than in H samples, and because the BV condition has about an order of magnitude more bacteria than does the H condition.

**Figure 3:**
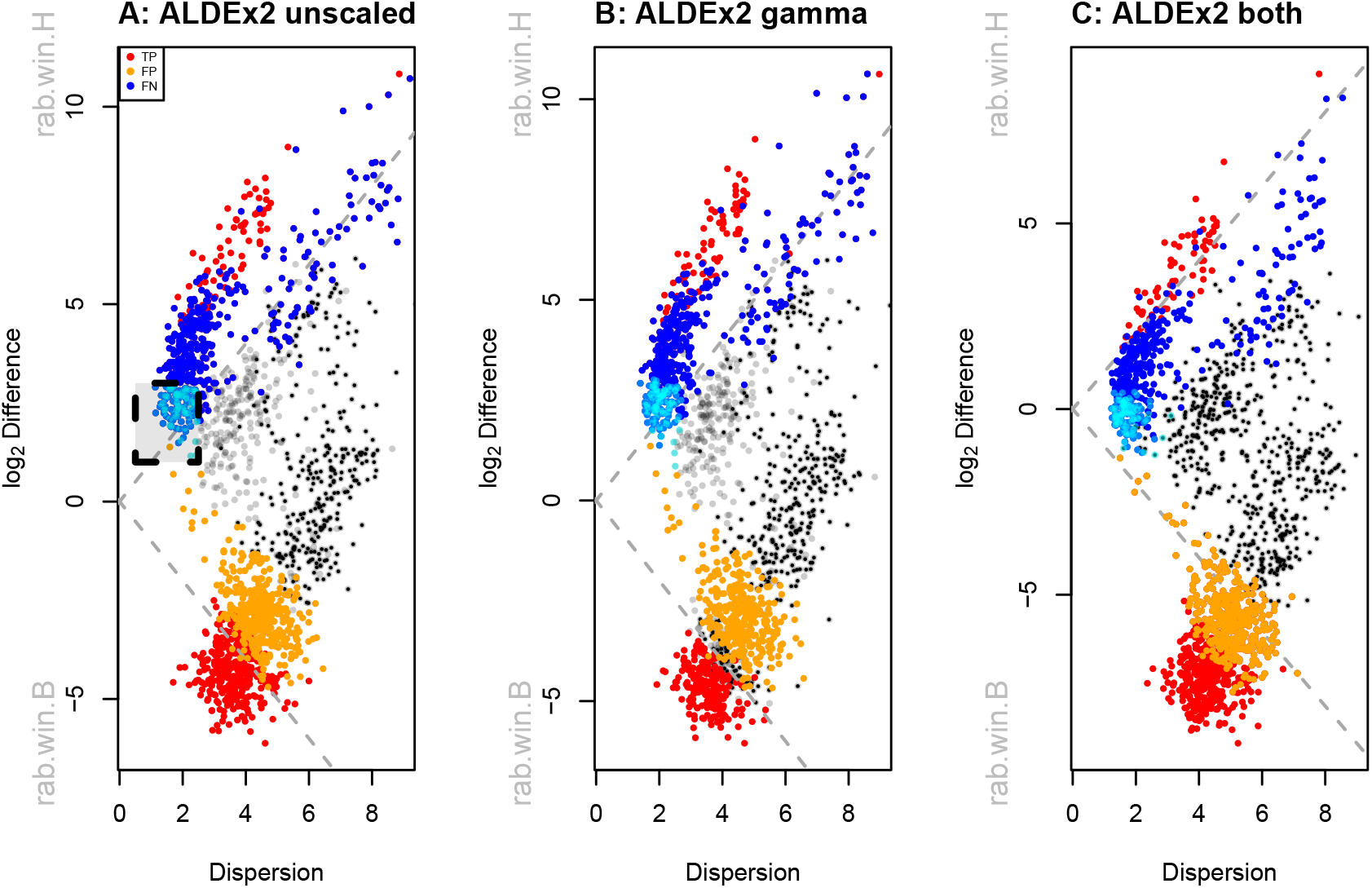
Analysis of vaginal transcriptome data aggregated at the Kegg Orthology (KO) functional level. Panel A shows an effect plot for the default analysis where the functions that are elevated in the healthy individuals have positive values and functions that are elevated in BV have negative values. Highlighed in the box are KOs that are almost exlusively housekeeping functions; these and are colored cyan. These housekeeping functions should be located on the midline of no difference. Panel B shows the same data scaled with *γ* = 0.5, which increase the minimum dispersion as before. Panel C shows the same data scaled with *γ* = 0.5 and a 0.14 fold difference in dispersion applied to the BV samples relative to the H samples. In these plots statistically significant (q-value < 0.01) functions in the informed model are in red, false positive functions are in blue, non-significant functions in black and false negative functions are in orange.

The naive scale model appears to be reflecting the bacterial load as observed by calculating the mean scale value for each group. Using a negligible scale value; i.e., *γ* = 1*e* − 3 exposes the naive scale estimate for samples in the @scaleSamps slot from the `aldex.clr` output. the naive scale estimate for the healthy group is 17.41 and for the BV group is 14.59 for a difference of 2.82. This is interpreted as the scale of the H group of samples being 7.06 greater than the BV group. This precise but incorrect estimate places the location of the housekeeping genes off the midline of no difference.

Applying the default scale model of *γ* = 0.5 increases the dispersion slightly but does not move the housekeeping functions toward the midline. This is as expected; the mean of the default scale model is based on the CLR normalization so no shift in location would be expected over the original ALDEx2 model. Nevertheless, about 30% of the housekeeping functions are no longer statistically significantly different. Note that this change is simple to conduct, has no additional computational complexity and requires only a slight modification for the analyst.

There are 101 functions with low dispersion that appear to be shared by both groups (boxed area in Figure 3A, and colored in cyan). Inspection shows that these largely correspond to core metabolic functions such as transcription, translation, ribosomal functions, glycolysis, replication, chaperones, etc (Supplementary file housekeeping.txt). The transcripts of many of these are commonly used as invariant reference sequences in wet lab experiments (48) and so are not be expected to contribute to differences in ecosystem behaviour. The average location of these should be centred on 0 difference to represent an internal reference set. However, without an informed scale model, the mean of these housekeeping functions is approximately located at +2.3.

We desire a scale model that approximately centres the housekeeping functions, because we expect housekeeping functions tp be nearly invariant; thus an appropriate scale in this dataset for functional analysis is likely closer to 0 than the naive estimate. One way to choose an appropriate value for *µ*_*n*_ is to use the `aldex.clr` function on only the presumed invariant functions setting *γ >* 0, and then accessing the @scaleSamps slot as before. Doing so suggests that the difference in scale should be about 14%. A second approach would be to identify the functions used as the denominator with the `denom=“lvha”` option (19) for the `aldex.clr` function, and then to use these values as before. This approach suggests a 5% difference in scale, and is potentially less subject to user interpretation.

For the purposes of this example, if we assume a 14% difference in scale, we can set *µ*_*i*_ = 1 and *µ*_*j*_ = 1.14 using the `makeScaleMatrix` function. This function uses a logNormal distribution to build a scale matrix given a user-specified mean difference between groups and uncertainty level. Applying a per-group relative differential scale of 0.14 moves the housekeeping functions close to the midline of no difference (Figure 3C, assuming 14% mean difference = -0.24, assuming a 5% mean difference = -0.34), and applying a gamma of 0.5 provides the same dispersion as in panel B of Figure 3. Note that now a significant number of functions are differentially up in BV that were formerly classed as not different without the full scale model (orange), or when only a default scale was applied. Inspection of the functions shows that these are largely missing from the *Lactobacillus* species and so should actually be captured as differentially abundant in the BV group. Supplementary Figure 4 shows that the using either the 5% or the 14% scale difference give imperceptibly different results suggesting that an informed scale model does not have to precisely estimate the scale difference to be useful. Nixon et al, (20) also found that multiple reasonable estimates for the *µ*_*n*_ part of the informed scale model were similarly useful in microbiome data.

Thus, applying an informed scale allows us to distinguish between both false positives (housekeeping functions in cyan, and others in blue) and false negatives (orange functions) even in a very difficult to analyze dataset. The remarkable improvements in biological interpretation afforded by an informed scale model, and the transferrability of it between sample cohorts of the same condition is outlined elsewhere (50). We suggest that the default scale model is sufficient when the data are approximately centred. However, an informed model is more appropriate with datasets are not well centred or when the investigator has prior information about the underlying biology.

## Discussion

Biological systems are both predictably variable and stochastic (49) and systems biology experiments show that there are transcripts with approximately constant concentrations in the cell and those with large variability under different growth conditions (23). Current measurement methods that rely on high throughput sequencing fail to capture all of the variation, particularly variation due to scale (3, 20). In the absence of external information (5, 6, 55) sequencing depth normalisation methods cannot recapture the scale information (5, 21), and can only normalize for the technical variation due to sequencing depth. Here we demonstrated that even approximate estimates of the true system scale and the uncertainty of measuring it can aid in the interpretation of RNA-sequencing experiments.

Nixon et al. (3) introduced the idea of explicitly modeling the scale of a HTS dataset, and showed how to incorporate these models in the analysis of microbiome and other datasets (20). They demonstrated that many tools commonly used to analyze HTS datasets had substantial Type 1 and Type 2 error rates, in line with recent findings by others (10, 12, 13). A version of ALDEx2 with the ability to include scale uncertainty was shown to be able to correct for the high Type 1 error rate for that tool, albeit with some loss of sensitivity. Finally, they showed that incorporating an informed scale model incorporating both location and scale uncertainty estimates could both control for Type 1 and Type 2 error rates (20).

Building and using a scale model thus has substantial benefits relative to the dual cutoff approach that is advocated for many gene expression experiments (14, 16). In particular, the dual cutoff approach has long been known to not control for Type 1 errors (17, 18), and the frequent lack of concordance between tools when benchmarked on transcriptomes(10, 12–14, 29, 56) and microbiomes (9, 11, 31, 40, 57, 58) suggests poor control of Type 2 errors as well (10, 13). Thus, incorporating a scale model during the analysis of HTS data promises the best of both worlds. A default scale model can control for Type 1 errors with minimal prior knowledge of the environment and this can be done with essentially no additional computational overhead. Furthermore, this work and previous (20) show that even minimal information about the underlying environment can be used to build a relatively robust informed scale model that controls for both Type 1 and 2 error rates.

In the analysis of HTS data it is often observed that larger datasets converge on the majority of parts being significantly different (3, 13, 14). Li et al. (13) conducted a permutation-based benchmarking study and found that widely used tools performed worse than simple Wilcoxon rank-sum tests in controlling the FDR when sample sizes became large. They suggested that the presence of outliers were one of the factors driving this observation. Brooks et al. (59) suggested that inappropriate choice of benchmarking methods are also a major contributing factor and that objective standards of truth are important. From the perspective of our work the disagreement between tools can be explained by the observation that different analytic approaches produce different parameter estimates for location or scale or for both. Thus, more data produces worse estimates because the additional data simply increases the precision of a flawed estimate (3, 60).

Scale simulation is now built into ALDEx2 (20) and here we suggest that there are two main root causes to common HTS data pathologies. The first contributing factor is the observed very low dispersion estimate for many features that is a by-product of some experimental designs and of normalization. In the Schurch et al. (14 dataset), the data were from single colonies derived from a single culture. Thus, it is more accurate to describe the 96 samples as wet-lab technical replicates rather than independent samples. This type of replication approach is standard in the molecular literature, and would be expected to result in the very low dispersion that is observed. Applying the default scale model with *γ* = 0.5 a large number of transcripts have their dispersion increased (Figure 1D), with the effect being largest for those with the lowest initial dispersion (Figure 2). Adding scale uncertainty results in modest number of transcripts, 205, being called significantly different as shown in the volcano plot in Figure 1B (red points). In addition, there is now a strong concordance between the difference and q-values. In hindsight, it is not obvious why the unscaled volcano plot shows such poor correspondence. We suggest that this is explained by random fluctuations of the many very low variance estimates and this is supported by the plots shown in Figure 2.

The second contributing factor is unacknowledged asymmetry in many datasets (19); i.e., different gene content or a directional change in the majority of features. In the case of asymmetry, the use of a user-specified scale model can be very useful for otherwise difficult-to-analyze datasets such as meta-transcriptomes and in-vitro selection datasets where the majority of features can change. We showed one such example for the metatranscriptome dataset in Figure 3. Here the dataset was highly asymmetrical. Incorporating differential scale on a per-group basis moves the mass of the housekeeping functions towards the midline of no difference and so affects both Type I and Type II error rates. We showed two ways of estimating the scale difference between groups and found that any reasonable estimate is an improvement over the naive approach and also over the default scale model. This is in line with the observations by Nixon et al (20) in a 16S rRNA gene sequencing dataset. It is also of note that in the case of true biological replicates (different individuals) that adding a modest amount of scale *γ* = 0.5 had little effect on the the difference between groups and on the dispersion. Thus, in this dataset the scale mis-specification was affecting mainly the location of the difference between groups. While we acknowledge that some prior information on which housekeeping transcripts should not be classed as differentially abundant is needed, we suggest that this information is widely available and is already used when performing the gold-standard quantitative PCR test of differential abundance (61, 62).

Beyond concerns of fidelity and rigor, scale models also enhance the reproducibility and transparency of HTS analyses. The addition of scale uncertainty essentially tests the model over a range of normalizations (3) and so can replace the consensus approach that has been proposed by some groups (11, 63) with no additional computational overhead. Thus, an advantage of incorporating scale is that analyses can be made much more robust such that actual or potential differences in scale can be tested and accounted for explicitly. While it is beyond the scope of the present article, we note that there are many ways of building scale models that enhance the interpretability of the parameters and assumptions and a detailed description of these points is describe elsewhere (3,).

In summary, we supply a toolkit that makes incorporating scale uncertainty and location information simple to incorporate for transcriptomes or indeed any type of HTS dataset. While the underlying scale of the system is generally inaccessible, the effect of scale on the analysis outcomes can be modelled and can help explain some of the underlying biology, and help to expose known issues with the analysis of HTS data. Adding scale information to the analysis allows for more robust inference because the features that are sensitive to scale can be identified and their impact on conclusions weighted accordingly. Additionally, the use of informed scale models permits difficult to analyze datasets to be examined in a robust and principled manner even when the majority of features are asymmetrically distributed or expressed (or both) in the groups (50). Thus, using and reporting scale uncertainty should become a standard practice in the analysis of HTS datasets.

