## supplementary figures for "Explicit Scale Simulation for analysis of RNA-sequencing with ALDEx2"

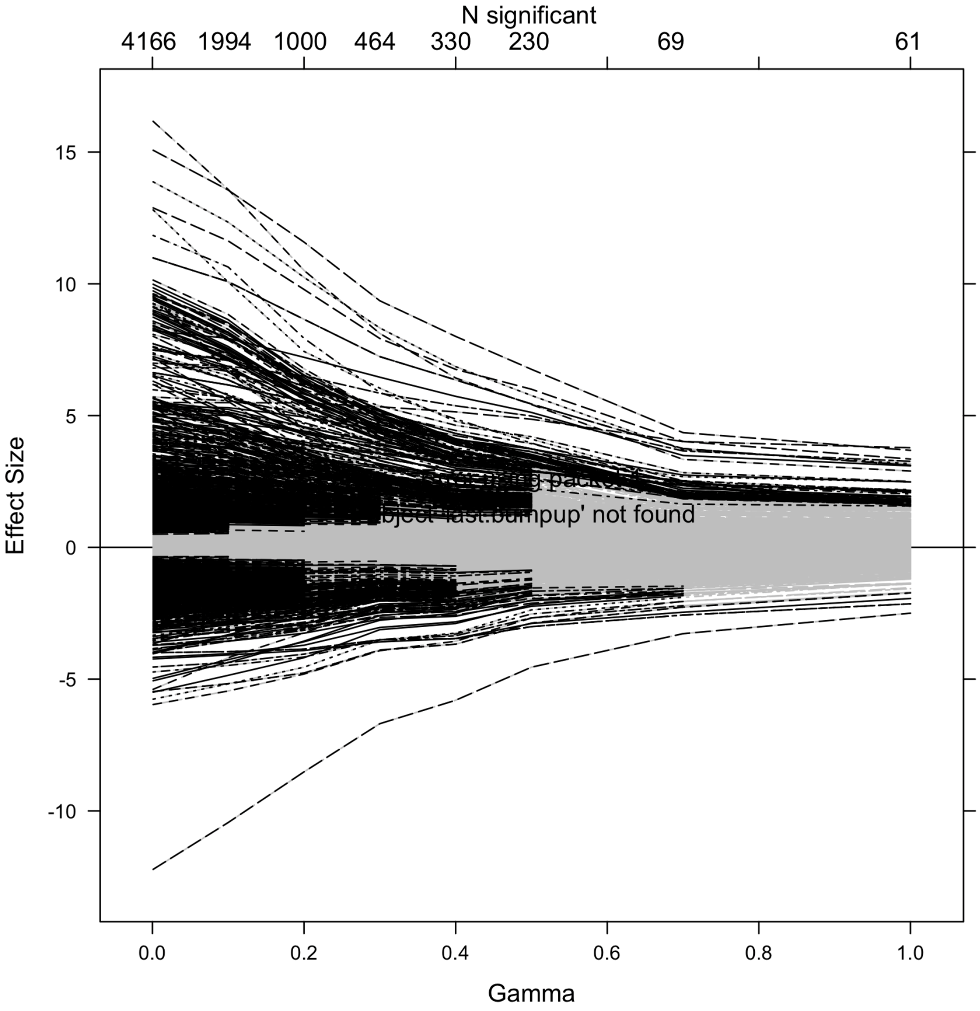


Figure 1: The aldex.senAnalysis() function was used to generate a dataset with $\gamma$ values of 1e-3, 0.1, 0.2, 0.3, 0.4,0.5, 0.7 and 1. The plotGamma() function was used to plot the result. Transcripts that are statistically significant are shown in black, and if not significant are in grey. The $\gamma$ values, standardized effect size and number of significant transcripts at each value are given on the axes.

One root cause of the large number of significant parts is the very low dispersion of transcripts. Figure 1 shows a graphical output from the texttt{aldex.scaleSim()} function with the yeast transcriptome dataset described in the text (1, 2). This allows us to examine which transcripts are sensitive to even minimal amounts of scale uncertainty using $\gamma = (1e-3, 0.1, 0.2, 0.3, 0.4, 0.5, 0.7, 1)$. Here it is obvious that even a negligible amount of scale uncertainty removes over half of the transcripts that were formerly significantly different, and all of these had very small effect sizes.

We recommend a minimum scale uncertainty of 0.5, and suggest that the texttt{aldex.senAnalysis()} function be run on all analyses. The individual `aldex()` outputs can be accessed as sequential entries in the list output, or the analysis as a whole can be plotted with the `plotGamma` function.

Figure 2 compares the significant transcripts using effect and volcano plots for the first two $\gamma$ parameters. We can see that the 5891 transcripts are excluded by even the smallest setting of $\gamma = 0.1$. All of these have very low difference between and include the majority of the transcripts with very low dispersion.

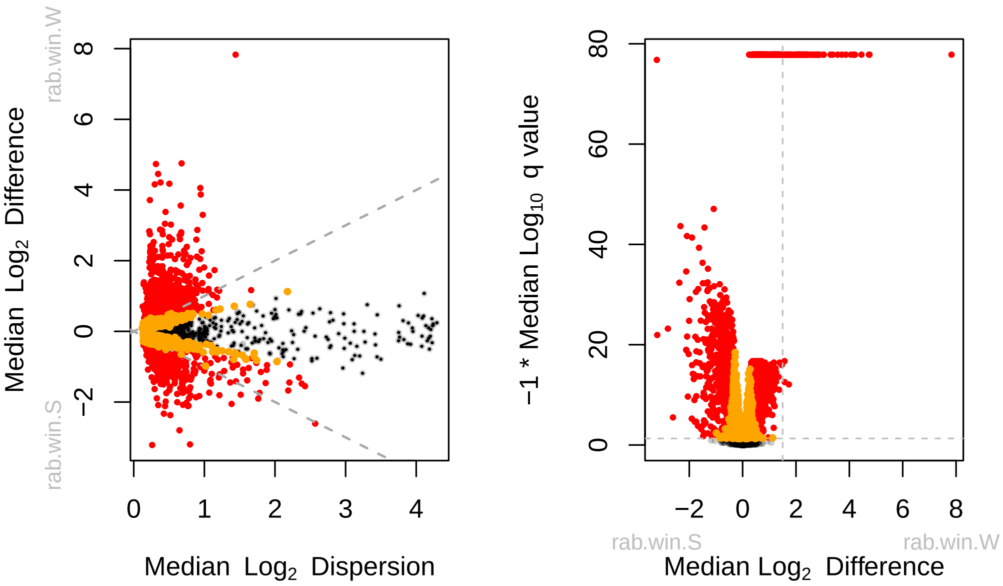


Figure 2: Effect and volcano plots showing the significant transcripts for the first two $\gamma$ values of 1e-3 and 0.1. Transcripts that are significantly different with a q-value <= 0.5 with both $\gamma$ values are in red, those significant with $\gamma = 0.1$ are in orange. Those that are not significant are in dark grey.

Figure 3 shows how dispersion behaves in linear and log space. The variance or dispersion always increases with increasing raw (or normalized) read count but decreases when measured on the log-ratio transformed data (3, 4), reaching a minimum at some mid-point of the distribution. This makes the counter-intuitive suggestion that genes with moderate expression have more predictable expression than genes with very high expression such as housekeeping genes. This is at odds with the known biology of cells where single cell counting of housekeeping transcripts shows that they are both highly expressed and have little intrinsic variation (5). Furthermore, the dispersion is exceeding small being, for many transcripts, almost negligible. To show this point more clearly, the majority of the transcripts in the lowest decile of dispersion indicated below the dashed grey line are statistically significantly different (75% with DESeq2, 69% with ALDEx2), suggesting that low dispersion estimates lead to many false positives. Indeed, benchmarking and comparison studies repeatedly show that the choice of normalization plays a role in which results are returned as significant (6–8). From the perspective of Nixon et al. (9), it is reasonable to conclude that some of these results may be due to the choice of normalization.

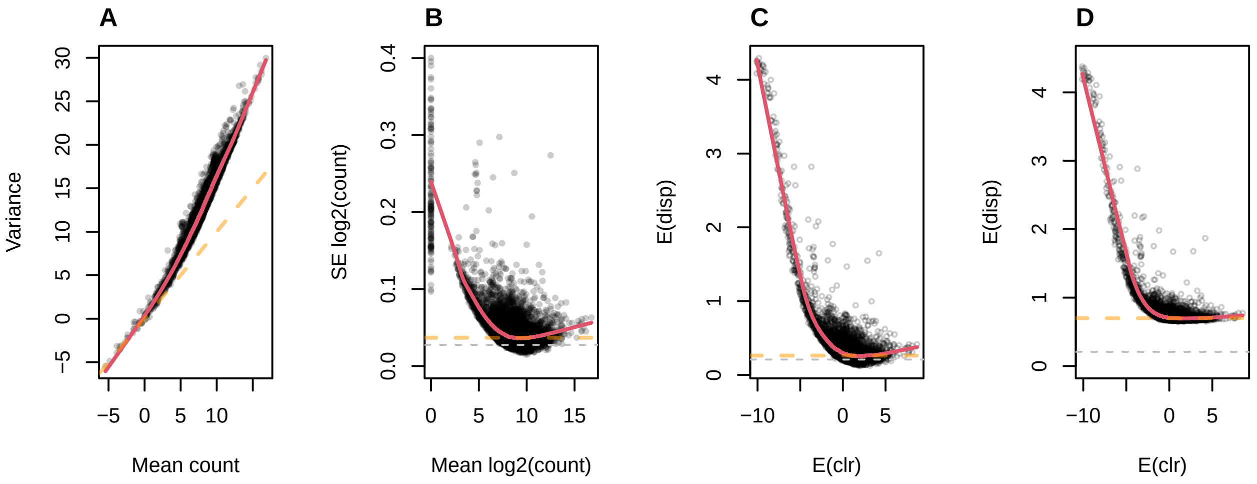


Figure 3: Plot of abundance v dispersion for the yeast transcriptome dataset as counts, as logarithms of counts, and as CLR values. Panel A shows that the data are over-dispersed relative to a Poisson distribution which is represented by the dashed line when plotted on a log-log scale. Panel B shows that the relationship between the mean and the dispersion calculated in DESeq2, here the standard error (SE) of the mean, is very different when the data are log-transformed first. Panel C shows the equivalent values calculated by ALDEx2 in which the expected CLR value for each transcipt are plotted vs. the expected dispersion. Panel D shows the output for ALDEx2 with $\gamma = 0.5$. The red line in each panel shows the LOESS line of fit to the mid-point of the distributions. In panels B and C the amount of dispersion reaches a minimum at moderate values. The dashed orange line in panel A is the line of equivalence, and in panel B and C is the minimum y value. The values below the dashed grey line in panels B and C represent those below the first decile of dispersion.

Figure 4 shows that the informed models with 5% or 14% difference in location between groups and $\gamma = 0.5$ provide nearly the same output for q-values, effect sizes and difference between groups (black). The line of identity is given as the grey diagonal. However, using the default CLR values to specify location are very different. In the q-value and effect plots, there is multiple populations of points that indicate the Type 1 and 2 errors that occur when the location is not specified properly. In the difference between groups plot, we see only a shift in the location that corresponds to moving the mass of the data points down by about 2.2 units (See Figure 3 in the main text)

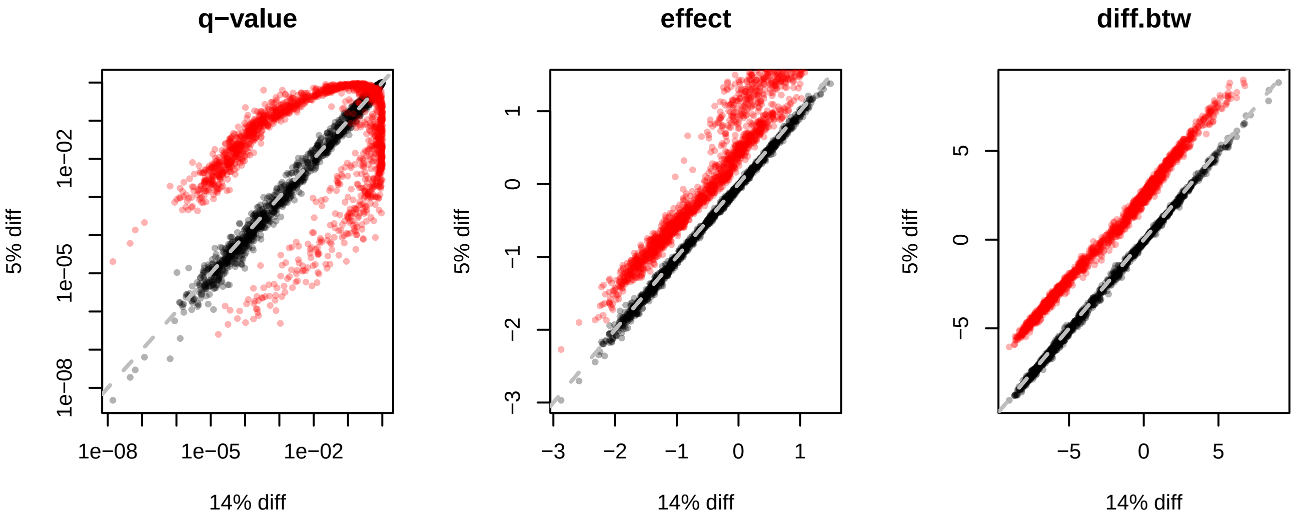


Figure 4: Plots showing the similarity of outputs with different scale parameters. The black points show that using either a an informed model with 5% or 14% difference in location has a minimal effect on either the q-values, the effect size or difference between groups. In red, the same values are plotted with the default model that uses a naive estimate of the location derived from the CLR.
